# Evolution of *Salmonella enterica* serotype Typhimurium driven by anthropogenic selection and niche adaptation

**DOI:** 10.1101/804674

**Authors:** Matt Bawn, Gaetan Thilliez, Mark Kirkwood, Nicole Wheeler, Liljana Petrovska, Timothy J. Dallman, Evelien M. Adriaenssens, Neil Hall, Robert A. Kingsley

**Affiliations:** Quadram Institute Biosciences, Norwich Research Park, Norwich, UK; Earlham Institute, Norwich Research Park, Norwich, UK; Centre for Genomic Pathogen Surveillance, Wellcome Sanger Institute, Cambridge, UK; Animal and Plant Health Agency, Addlestone, UK; Gastrointestinal Bacteria Reference Unit, National Infection Service, Public Health England, London, UK; University of East Anglia, Norwich, UK

## Abstract

*Salmonella enterica* serotype Typhimurium (*S*. Typhimurium) is a leading cause of gastroenteritis and disseminated disease worldwide. Two *S*. Typhimurium strains (SL1344 and ATCC14028) are widely used to study host-pathogen interactions, yet genotypic variation results in strains with diverse host range, pathogenicity and risk to food safety. A robust fully parsimonious phylogenetic tree constructed from recombination purged variation in the whole genome sequence of 131 diverse strains of *S*. Typhimurium revealed population structure composed of two high order clades (α and β) and multiple subclades on extended internal branches, that exhibited distinct signatures of host adaptation and anthropogenic selection. Clade α contained a number of subclades composed of strains from well characterized epidemics in domesticated animals, while clade β predominantly contained subclades associated with wild avian species, with the notable exception of a subclade containing the DT204/49 complex. The contrasting epidemiology of α and β strains was reflected in a distinct distribution of antimicrobial resistance (AMR) genes, accumulation of hypothetically disrupted coding sequences (HDCS), and signatures of functional diversification associated with invasiveness of host adapted serotypes. Gene flux was predominantly driven by acquisition, loss or recombination of prophage. The acquisition of large genetic islands (SGI-1 and 4) was limited to two recent pandemic clones (DT104 and monophasic *S*. Typhimurium ST34) in clade α. Together, our data are consistent with the view that a broad host range common ancestor of *S*. Typhimurium diversified with clade α lineages remained largely associated with multiple domesticated animal species, while clade β spawned multiple lineages that underwent diversifying selection associated with adaptation to various niches, predominantly in wild avian species.

## Introduction

Bacteria of the genus *Salmonella* are a common cause of foodborne disease. Most of the approximately 2500 serovars cause gastroenteritis in humans and other animals, while some have evolved host adaptation associated with extra intestinal disseminated infections in specific host species ^1^. For example, *Salmonella enterica* serotype Typhimurium (*S*. Typhimurium) and *S*. Enteritidis circulate in multiple vertebrate host species and cause food borne infections in the human population resulting in an estimated 75 million cases and 27 thousand deaths from gastroenteritis worldwide ^2^. *S*. Typhi and *S*. Paratyphi A circulate exclusively in the human population and cause an estimated 2.5 million infections resulting in 65 thousand deaths each year as a result of the disseminated disease typhoid and paratyphoid disease ^2^. Similarly, other serotypes evolved host adaptation to specific non-human host species, such as *S*. Gallinarum with poultry, *S*. Dublin with cattle, and *S*. Choleraesuis with pigs, where they are associated with disseminated infections ^1^.

Although *S*. Typhimurium is considered to be a broad host range serotype, the epidemiological record of *S*. Typhimurium phage types identifies several *S*. Typhimurium pathovariants with distinct host range, pathogenicity and risk to food safety ^3, 4^. The pathovariant commonly associated with this serotype, has a broad host range and is associated with gastroenteritis in the human population. Such broad host range strains of *S*. Typhimurium account for the majority of those isolated by public health surveillance in England, presumably because they are common in many species of livestock and poultry, the primary zoonotic reservoir for human infections ^5^. The epidemiological record of this pathovariant is characterised by successive waves of dominant clones identified historically by their phage type, that account for up to 60% of all human infections for several years, before being replaced by a subsequent strains ^6^. Dominant clonal groups have been characterized by strains of phage types DT9, DT204/49 complex, DT104, and the current monophasic *S*. Typhimurium (*S*. 4,[5],12:i:-) sequence type 34 (ST34), since around the middle of the last century ^7–10^. In contrast, some phage types are common in clonal groups typically associated with a restricted host range, and in some cases altered pathogenicity. For example, clonal groups of *S*. Typhimurium DT8, DT2 and DT56 circulate in populations of ducks, pigeon, and passerine birds, respectively, and only rarely cause gastroenteritis in the human population ^11–13^. Also, specific clonal groups of *S*. Typhimurium ST313 are associated with disseminated disease (invasive non-typhoidal *Salmonella*, iNTS) in sub-Saharan Africa^14, 15^.

In this study we report the population structure, gene flux, recombination and signatures of functional diversification in the whole genome sequence of 131 strains of *S*. Typhimurium with well characterised epidemiology. To assist in our analysis, we also report high quality complete and closed whole genome sequence of six additional reference genomes, representing diversity within the population structure not represented by previously reported sequence.

## Results

### Population structure of *S*. Typhimurium consists of two high-order clades containing strains with distinct epidemiology

Variant sites (38739 SNPs) in the core genome sequence of 134 *S*. Typhimurium strains representing commonly isolated phage types revealed two diverse phylogroups composed of three strains of ST36 that clustered separately from the remaining 131 Typhimurium isolates (Supplementary Figure 1). The two clusters were more similar to one another than any other serotype, including the most closely related serotype, *S*. Heidelberg. Since the majority of *S*. Typhimurium formed a large number of relatively tightly clustered isolates, we focussed on the analysis of the population structure and evolution of this phylogroup. A phylogenetic tree constructed using variant sites (8382 SNPs) in the core genome sequence of the 131 *S*. Typhimurium strains and rooted with *S.* Heidelberg, revealed a ‘star’ topology with relatively long internal branches extending from a hypothetical common ancestor, and diversification at the terminal branches (Figure 1). The population structure determined using a three-level hierarchical Bayesian approach ^16^ resolved *S*. Typhimurium into two major clades, designated α and β, and third-level clades (α 8-18 and β 1-7) that corresponded to known epidemic clades. Clade β was defined by an internal branch defined by approximately 100 core genome SNPs that originated from a common ancestor of the basal clade α.

**Figure 1.**
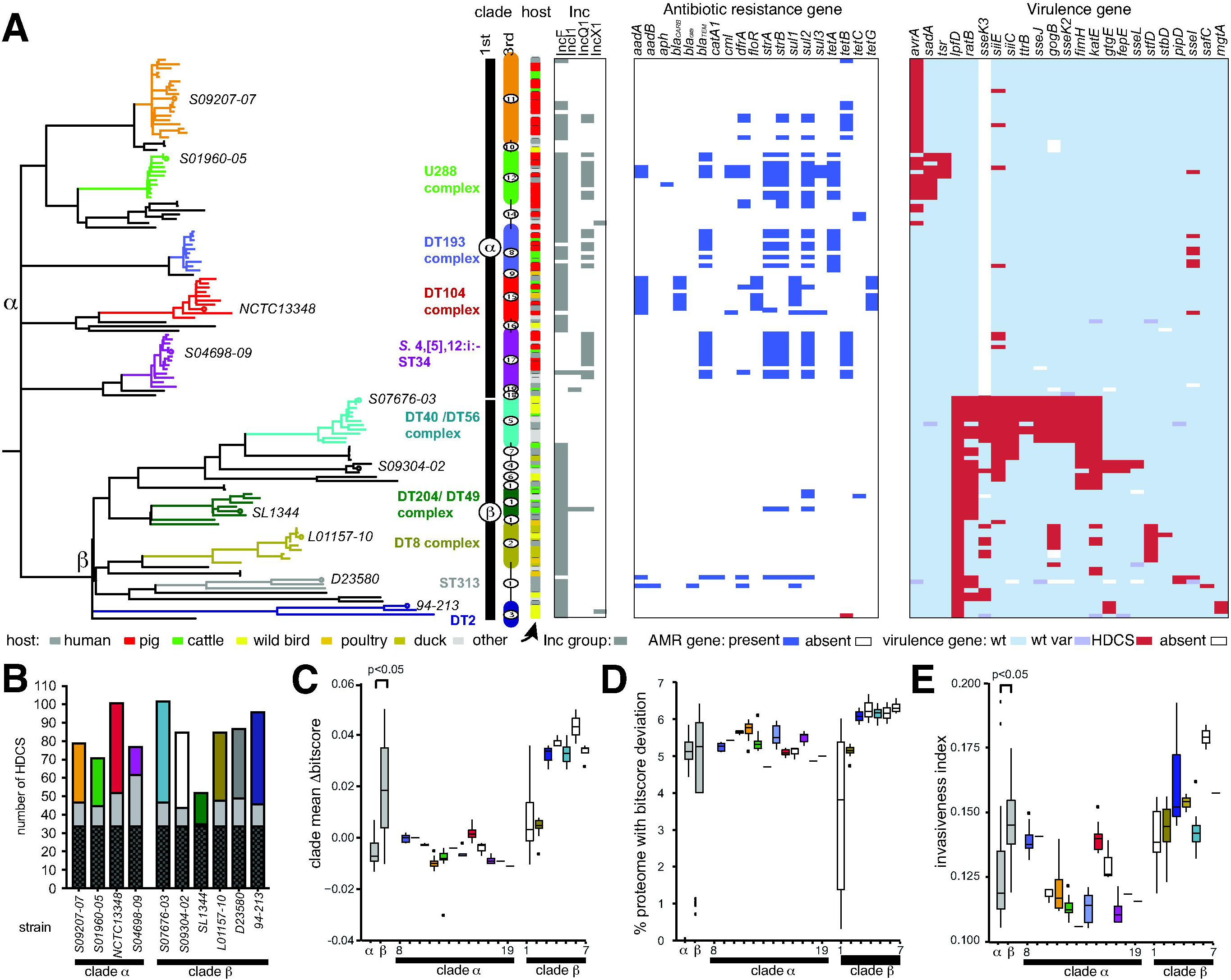
Phylogenetic relationship of the ST19 *Salmonella* Typhimurium phylogroup. (A) Maximum likelihood phylogenetic tree and based on sequence variation (SNPs) in the core genome with reference to S. Typhimurium strain SL1344. The root was identified using *S.* Heidelberg (accession number NC_011083.1) as the outgroup. 1^st^ (α and β) and 3^rd^ (α11-19 and β1-7) are indicated (vertical bars). Phage type complexes associated with the third-level clusters are indicated (bold type) colour coded with the lineages and representative strains from third level clusters (italicized type). The source of each isolate in the tree is indicated by filled boxes colour coded as indicated in the inset key (arrow). The presence of replicon sequence (grey box), antimicrobial resistance genes (blue box) and hypothetically disrupted coding sequence (HDCS) of virulence related genes (red box) in short read sequence data are indicated. (B) Bars indicate the number of ancestral (black), phage or insertion sequence elements (grey), chromosomal gene (colour coded with lineages in Figure 1A) HDCS in the genome of representative strains from each third level clade. (C) Box plots indicate the mean dbitscore (DBS: bitscore SL1344 – test strain bitscore) of proteomes in third level clades. (D) Box plot indicates the percentage of the proteome of the proteome of isolates from each third level clade with a non-zero bitscore (bitscore SL1344 – test strain bitscore >0 or <0) as an estimate of function divergence. (E) Box plots indicate the mean invasiveness index per genome, the fraction of random forest decision trees voting for an invasiveness phenotype based on training on the DBS of a subset of the proteome of ten gastrointestinal and extraintestinal pathovar serotypes.

Despite relatively few SNPs distinguishing clade α and β, these clades exhibited distinct epidemiology characterised by association predominantly with livestock (clade α) or avian species, including wild species (clade β). Strikingly all pig isolates from our sampling were located in clade α. Cattle isolates were in both first order clades (11 in clade α 7 in clade β), but in clade β they had a relatively limited distribution with five of the isolates from a subclade containing the DT204/49 complex of strains associated with a cattle associated epidemic in the 1970’s ^8^. Clade α contained strains from several previously described epidemics in livestock animal species, including in pigs (α12, U288)^17^, two clades associated with recent pandemic clonal groups associated with pigs, cattle and poultry (α17, monophasic Typhimurium ST34 and α15, DT104) ^18–20^, and potentially epidemic clades not previously described in the literature, consisting of isolates from pigs, cattle and poultry (α8 and α11). Clade-β was characterised by many long internal branches, indicative of a relatively high level of sequence divergence, relative to those in Clade-α. In contrast, clade-β contained several third-level clades previously described as host-adapted, particularly for avian species such as passerine birds (β5, DT56), duck (β2, DT8) and pigeon (β3 DT2) ^21^, and ST313 that includes two sub-clades specifically associated with disseminated disease in sub-Saharan Africa ^22^.

### Antimicrobial resistance genes and plasmid replicons are associated with strains associated with livestock

The presence of multiple third-level clades associated with recent livestock associated epidemic strains in the first level clade α and the relative paucity in clade β suggested that they may be under differential anthropogenic selection pressure. A key anthropogenic selection pressure on microbial populations circulating in livestock is the widespread use of antimicrobial drugs in animal husbandry. Consistent with their distinct epidemiology, antimicrobial resistance (AMR) genes were common in clades α (mean of 2.7 per strain) and relatively rare in β (mean 0.38 per strain) (Figure 1A and Supplementary Table 2). Indeed, AMR genes in clade-β were restricted to strains from DT204/49 complex known to be associated with cattle ^8^ and the ST313 associated with disseminated disease in sub-Saharan Africa commonly treated with antibiotics ^22^. AMR genes are commonly present on plasmids and we therefore determined the presence of plasmid replicon sequence in short read sequence data from the 131 strains in clades α and β. The IncF replicon corresponding to the presence of the virulence plasmid pSLT ^23^ was widespread in *S*. Typhimurium, but notably absent from a number of third-level clades including β5 (DT56), α11 and α17 (monophasic *S*. Typhimurium ST34). The IncQ1 plasmid replicon, previously associated with antibiotic resistance ^24^, was also widespread, particularly in clade α, associated with domesticated animal species.

### Distinct patterns of genome degradation and signatures of functional divergence and invasiveness in clades α and β

Hypothetically disrupted coding sequences (HDCS) due to frameshift mutations or premature stop codons were determined in high quality finished and closed genome sequence of 11 representative strains from major subclades (Figure 1B and Supplementary Table 3). Representative strains of clade α generally contained fewer HDCS than those from clade β, with the exception of NCTC13348 (DT104) and SL1344 (DT204/49) that had atypically high and low numbers of HDCS, respectively (Figure 1B). However, none of the HDCS in NCTC13348 were in genes previously been implicated in pathogenesis, while SL1344 contained two virulence gene HDCS (*lpfD* and *ratB*) (Figure 1A). In general, clade α strains had 0-3 HDCS in virulence genes (mean 0.8 SD 0.9), while clade β was characterised by multiple lineage containing three or more virulence gene HDCS (mean 5.0 SD 3.2) (Figure 1A and Supplementary Table 4). Isolates in clade β5 (DT56, passerine bird associated ^13^) contained up to eleven virulence gene HDCS (*lpfD*, *ratB*, *sseK3*, *siiE*, *siiC*, *ttrB*, *sseJ*, *gogB*, *sseK2*, *fimH* and *katE*). The greatest number of HDCS in clade α were observed in α12 (U288, possibly pig adapted), in which three virulence genes were affected (*avrA*, *sadA* and *tsr*). *LpfD* was found to be the only HDCS (Supplementary Table 3) that segregated between clade-α and clade-β. A 10-nucleotide deletion causing a frameshift mutation in *lpfD* resulting in a truncation approximately half way into the protein in all isolates of clade-β.

To quantify the relative level of functional divergence in the proteome of isolates in each clade we used a profile hidden Markov Model approach, delta-bitscore (DBS profiling) ^25^ (Figure 1C and Supplementary Table 5). The method assigned a value (bitscore) to peptides of the proteome that indicated how well each sequence fitted the HMM. We determined the difference in bitscore of the proteome of each isolate relative to that of *S*. Typhimurium strain SL1344 (DBS = bitscore SL1344 proteome - bitscore test proteome). A greater DBS is therefore indicatives of excess of polymorphisms that potentially alter protein function, and most likely a loss of function as it indicates divergence from the profile HMM. Mean DBS was significantly greater (p<0.05, Wilcoxon test) for proteomes of strains in clade-β compared with clade-α (Figure 1C). In general, third-level clades in clade-α exhibited DBS of approximately zero, consistent with limited functional divergence. Notably, despite considerable numbers of HDCS in strain NCTC13348 (α15, DT104), DBS was only moderately elevated in this clade. Proteomes of strains in clade β exhibited mean DBS of 0.03 and above with the exception of clades β1 and β2. The proportion of the proteome with any deviation in DBS was also greater in clade-β than clade-α (Figure 1D).

We also used a machine learning approach to predict the ability of strains to cause extraintestinal disease based on the DBS of 196 proteins that were recently determined as the most predictive of the invasiveness phenotype in 13 serotypes (six extra-intestinal pathovars and seven gastrointestinal pathovars) of *S. enterica* subspecies I ^26, 27^ (Figure 1E). The invasiveness index metric is the fraction of decision trees in a random forest algorithm that vote for an invasive phenotype. The invasiveness index was significantly greater (p<0.05, Wilcoxon test) in clade-β than clade-α, consistent with the epidemiology and pathogenicity of the isolates located in these clades ^3, 4^.

### The clade-specific accessory genome is largely driven by acquisition of prophage genes and integrative elements

A pangenome analysis of *S*. Typhimurium (excluding the ST36 phylogroup) identified 9167 total gene families. The core genome (present in 99-100% of strains) was 3672 genes, soft core genome (95-99%) 388 genes. Shell genome (15-95%) 792 genes, and cloud genes (0-15%) 4315 genes (Figure 2A and 2B). We defined gene families of the accessory genome as non-prophage chromosomal, prophage, plasmid and undefined, based on their location and annotation in the complete and closed genomes of eleven reference strains phylogenetically distributed across S. Typhimurium (Table 1). Gene families not present in reference genomes were classified as ‘undefined’. The accessory genome was defined as genes present in 95% or fewer of isolates and thus represents the major source of genetic variation between strains.

**Figure 2.**
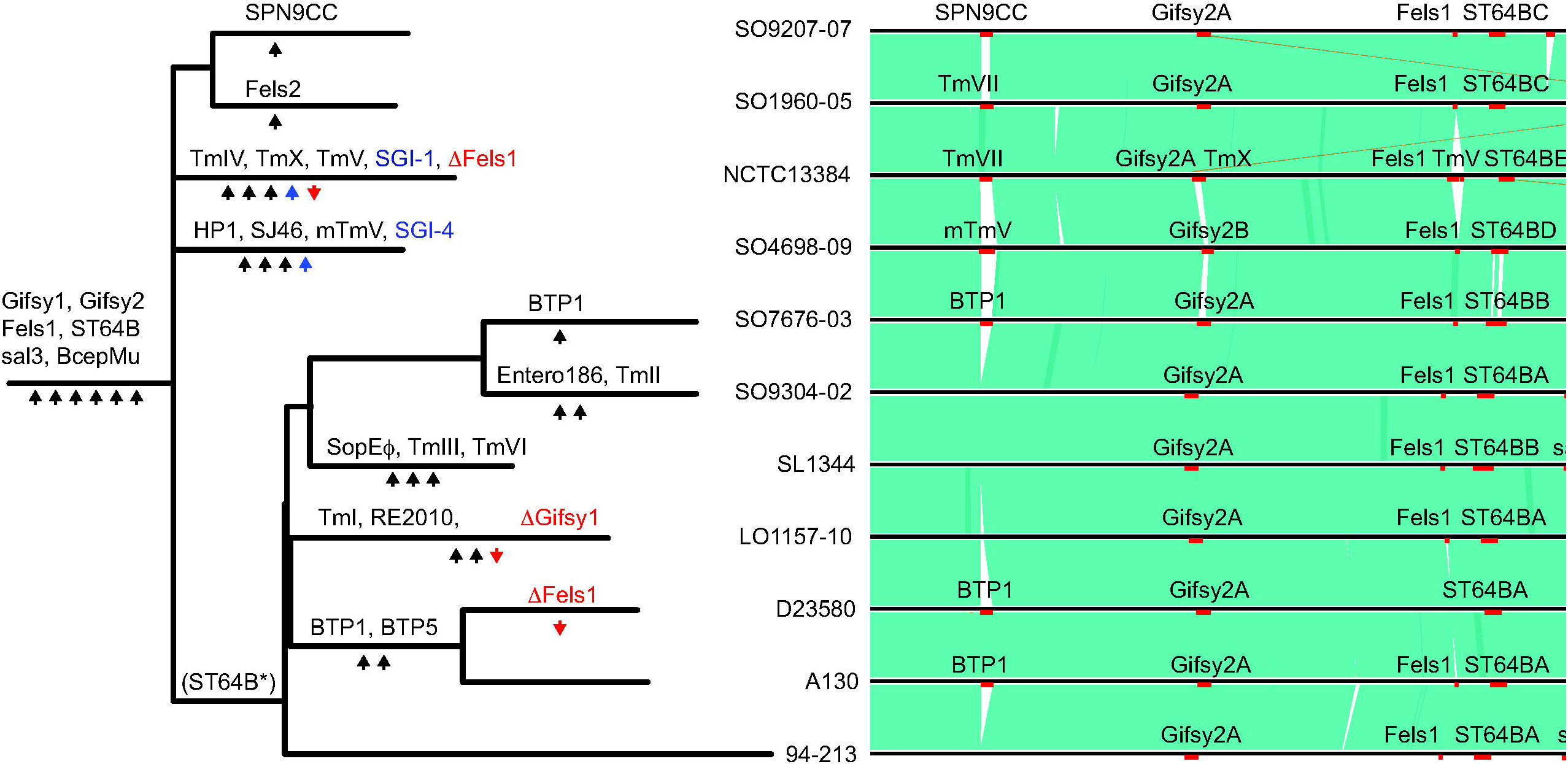
The pan genome of 131 *S*. Typhimurium isolates. Gene families were identified based on sequence alignment with a cut off of 90% sequence identity and assigned to non-prophage chromosomal (red), prophage (green), plasmid (blue), or undefined (grey), based on their genome context in eleven annotated reference genomes from each third level clade. (A) Number of genome families in the core, softcore, shell and cloud components of the pangenome. (B) Number of genome families of each pan genome component in isolates from each *S*. Typhimurium third level clade. (C) Accessory genome (shell and cloud) in each isolate. Gene families present in more than 130 or less than 5 strains were excluded. Maximum Likelihood tree based on variation (SNPs) in the core genome with reference to S. Typhimurium SL1344. Third-level clades are indicated in colour coded in common with the phylogeny vertical bars.

**Figure 3.**
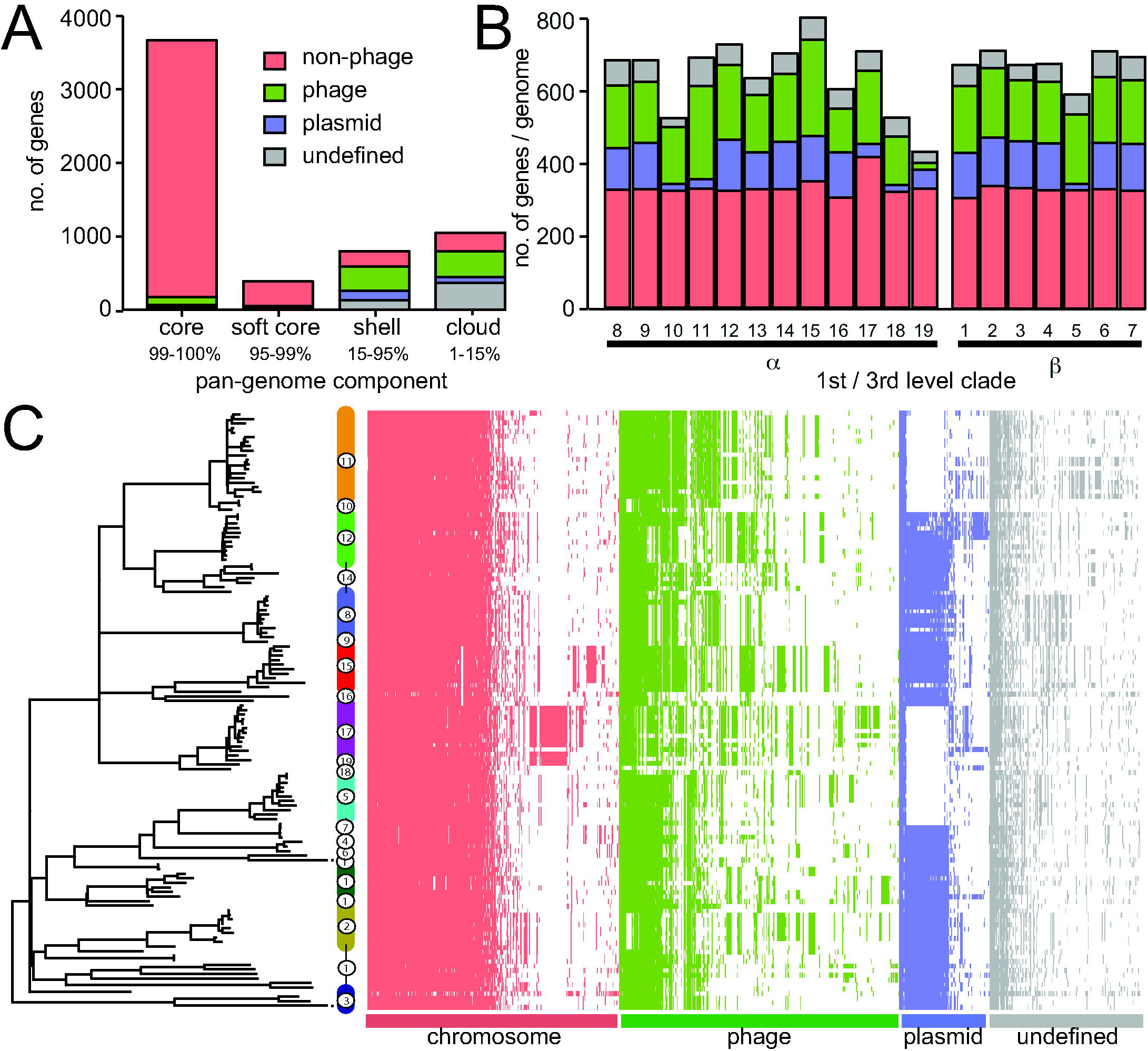
Genome alignment and phylogenetic relationship of complete and closed reference strains of S. Typhimurium or used in this study. Sequence with >90% nucleotide sequence identity are indicated where this is direct alignment (green) or reverse and complement (red). The location of prophage sequence (red bars) or integrative elements (blue bars) are indicated. A maximum likelihood tree based on sequence variation (SNPs) in the core genome with reference to S. Typhimurium strain SL1344 (left) is annotated with the most likely order of acquisition (black arrow) or loss (red arrow) of prophage and integrative elements, based on the principle of parsimony.

**Table 1.**
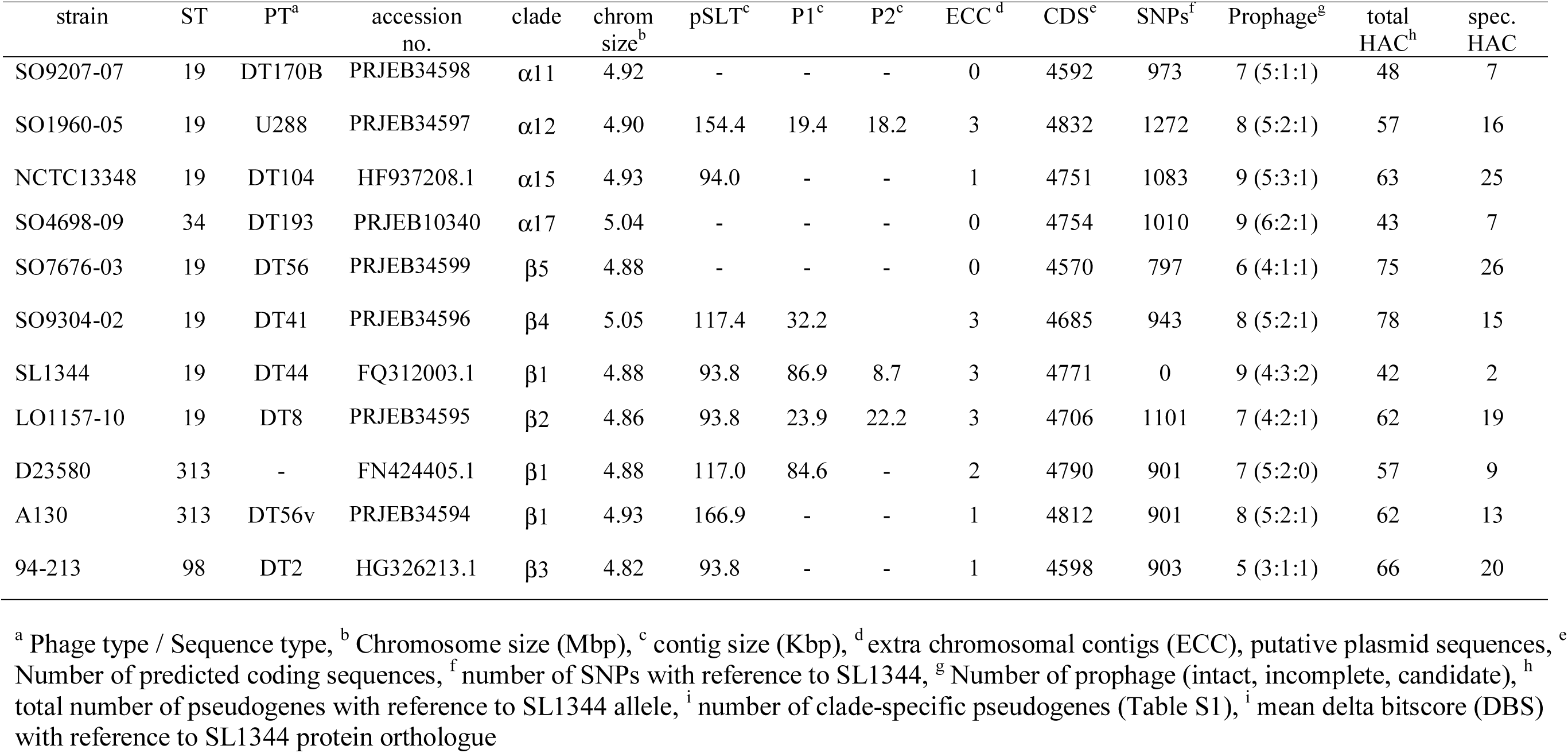
Characteristics of complete and closed whole genome sequence of *S*. Typhimurium reference strains.

| strain | ST | PT<sup>a</sup> | accession no. | clade | chrom size<sup>b</sup> | pSLT<sup>c</sup> | P1<sup>c</sup> | P2<sup>c</sup> | ECC<sup>d</sup> | CDS<sup>e</sup> | SNPs<sup>f</sup> | Prophage<sup>g</sup> | total HAC<sup>h</sup> | spec. HAC |
| --- | --- | --- | --- | --- | --- | --- | --- | --- | --- | --- | --- | --- | --- | --- |
| SO9207-07 | 19 | DT170B | PRJEB34598 | α11 | 4.92 | - | - | - | 0 | 4592 | 973 | 7 (5:1:1) | 48 | 7 |
| SO1960-05 | 19 | U288 | PRJEB34597 | α12 | 4.90 | 154.4 | 19.4 | 18.2 | 3 | 4832 | 1272 | 8 (5:2:1) | 57 | 16 |
| NCTC13348 | 19 | DT104 | HF937208.1 | α15 | 4.93 | 94.0 | - | - | 1 | 4751 | 1083 | 9 (5:3:1) | 63 | 25 |
| SO4698-09 | 34 | DT193 | PRJEB10340 | α17 | 5.04 | - | - | - | 0 | 4754 | 1010 | 9 (6:2:1) | 43 | 7 |
| SO7676-03 | 19 | DT56 | PRJEB34599 | β5 | 4.88 | - | - | - | 0 | 4570 | 797 | 6 (4:1:1) | 75 | 26 |
| SO9304-02 | 19 | DT41 | PRJEB34596 | β4 | 5.05 | 117.4 | 32.2 | | 3 | 4685 | 943 | 8 (5:2:1) | 78 | 15 |
| SL1344 | 19 | DT44 | FQ312003.1 | β1 | 4.88 | 93.8 | 86.9 | 8.7 | 3 | 4771 | 0 | 9 (4:3:2) | 42 | 2 |
| LO1157-10 | 19 | DT8 | PRJEB34595 | β2 | 4.86 | 93.8 | 23.9 | 22.2 | 3 | 4706 | 1101 | 7 (4:2:1) | 62 | 19 |
| D23580 | 313 | - | FN424405.1 | β1 | 4.88 | 117.0 | 84.6 | - | 2 | 4790 | 901 | 7 (5:2:0) | 57 | 9 |
| A130 | 313 | DT56v | PRJEB34594 | β1 | 4.93 | 166.9 | - | - | 1 | 4812 | 901 | 8 (5:2:1) | 62 | 13 |
| 94-213 | 98 | DT2 | HG326213.1 | β3 | 4.82 | 93.8 | - | - | 1 | 4598 | 903 | 5 (3:1:1) | 66 | 20 |

<sup>a</sup> Phage type / Sequence type, <sup>b</sup> Chromosome size (Mbp), <sup>c</sup> contig size (Kbp), <sup>d</sup> extra chromosomal contigs (ECC), putative plasmid sequences, <sup>e</sup> Number of predicted coding sequences, <sup>f</sup> number of SNPs with reference to SL1344, <sup>g</sup> Number of prophage (intact, incomplete, candidate), <sup>h</sup> total number of pseudogenes with reference to SL1344 allele, <sup>i</sup> number of clade-specific pseudogenes (Table S1), <sup>i</sup> mean delta bitscore (DBS) with reference to SL1344 protein orthologue

Some gene families exhibited a distinct distribution in clade α or β, or within individual third-level clades of clade α or β (Supplementary Figure 2). Four non-phage chromosomal genes were specifically associated with clade-β strains, STM0038 a putative arylsulfatase, *tdcE* encoding a pyruvate formate lyase 4, *aceF* acetyltransferase and *dinI* a DNA damage inducible protein. A series of plasmid genes in clade β2 corresponded to a region of p2 seen in LO1157-10. This region is likely a transposon as it contains an IS200 transposase and an integrase. Also, a number of prophage-associated genes were present throughout clade-β due to apparent recombination in the ST64B prophage. The rate of gene flux in clade α and clade β was determined by computing the number of accessory genes as a function of SNPs in each clade. By this measure, the rate of gene flux was nearly twice as high in clade α compared to clade β (Supplementary Figure 3).

Generally, gene flux in the non-prophage and non-plasmids gene families that were specific to individual third level clades was limited to individual genes or small blocks of genes (Supplementary Figure 2). The exception was two large genetic islands in clades α15 (DT104 complex) and α17 (monophasic *S*. Typhimurium ST34) corresponding to the presence of SGI1 ^28^ and SGI4 ^29^. In addition, a chromosomal block of genes in clade β2 corresponding to an insertion at the Thr-tRNA at 368274 containing a series of hypothetical proteins and a gene with similarity to the *trbL* gene involved in conjugal transfer (A0A3R0DZN6) and a site-specific integrase. Some of these genes were also present in isolates in clades β1 and β3. The greatest contribution to third level clade specific gene families was in the those with predicted functions in prophage (Figure 2C and Supplementary Figure 2).

### Extant prophage repertoire is the result of recombination and infrequent loss of ancestral elements and acquisition of new phage

In order to investigate the flux of prophage genes resulting in clade-specific repertoires, we identified prophage in eleven complete and closed reference genomes of *S*. Typhimurium sequences. A total of 83 complete or partial prophage elements were identified in the eleven reference genomes (Figure 4). Prophage were present at twelve variably occupied chromosomal loci and the number per strain ranged from five in DT2 (strain 94-213) to nine in monophasic *S*. Typhimurium ST34 (strain SO4698-09) (Table 1). Clustering of gene families in the prophage pangenome identified 23 prophage, although in some cases blocks of genes were replaced resulted in mosaic prophage for example “*Salmonella* virus ST64BX” and “*Salmonella* virus Gifsy1X” (supplementary table 6), and the definition of families of prophage with a high confidence was consequently problematic (Figure 4). Thirteen prophage elements encoded at least one identifiable cargo gene, capable of modifying the characteristics of the host bacterial strain, including eleven genes previously implicated in virulence (Supplementary Table 3). Ten prophage families contained no recognisable cargo genes. The evolutionary history of prophage acquisition and loss was reconstructed based on principles of maximum parsimony. Six prophage (*Salmonella* viruses “BcepMuX”, “Gifsy1X”, “Gifsy2X”, “Fels1X”, “ST64BX” and “sal3X”, hereafter referred to as BcepMu, Gifsy1, Gifsy2, Fels1, ST64B and sal3) (supplementary table 6), that together accounted for 61 of the prophage in these genomes, were most likely present in the common ancestor of *S*. Typhimurium. Loss of two of these ancestral prophage by three isolates (NCTC13348, L01157-10 and D23580) represented the only evidence for decrease in prophage repertoire in the dataset.

**Figure 4.**
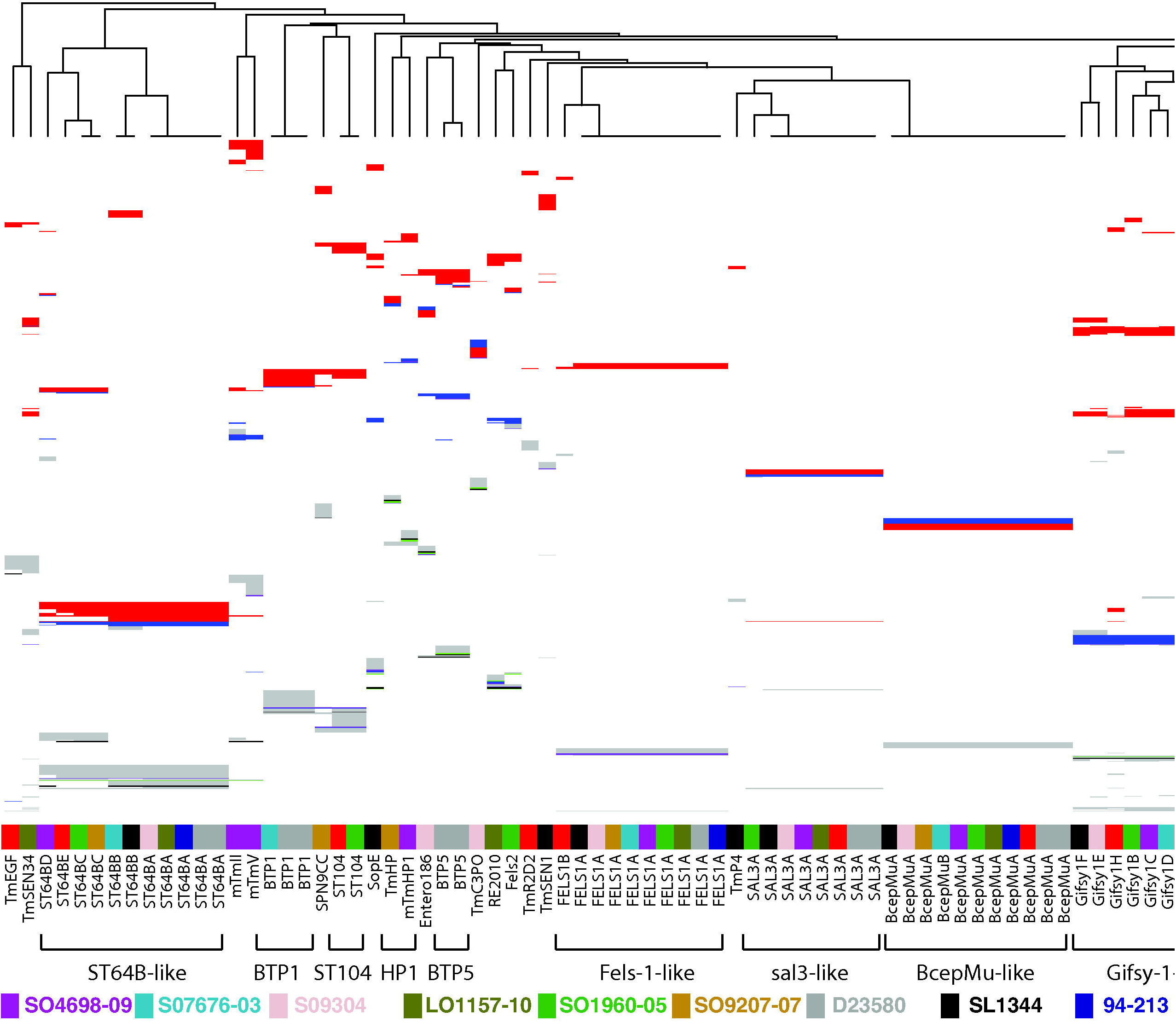
Clustering of prophage genes based on sequence identity indicates related families and potential recombination. Genes from all prophage identified in complete and closed whole genome sequence of eleven reference strains of *S*. Typhimurium were assigned to families based on sequence identity (>90% identity). Prophage genes (columns) were clustered to identify related prophage. The presence of a gene is indicated with a box predicted function based on *in silico* annotation are colour coded based on annotation, terminase (black), capsid (green), recombinase/integrase (purple), tail fibre (blue), other phage associated (red), and hypothetical protein (grey). A cladogram showing the relationship of prophage is based on the pattern of gene presence or absence is indicated (top).

A total of 22 additional prophage had a limited distribution within *S*. Typhimurium strains, present in three or fewer genomes *Salmonella* viruses (“TmEGF”, “TmSEN34”, “mTmII”, “mTmV”, BTP1, “TmST104”, “SPN9CC”, Fels2, “TmHP1/mTmHP1”, BTP5, “TmC3PO”, “RE2010”, “TmR2D2” and “TmSEN1”) (Table 2 and supplementary table 6) and are therefore likely to have been acquired during the evolution of *S*. Typhimurium (Figure 4). *Salmonella* virus BTP1 that was reported to be specific to the ST313 strains associated with epidemics of invasive NTS disease in sub-Saharan Africa ^30^, was also present in strain SO7676-03, a strain in clade β5 (DT56 complex) adapted to circulation in wild bird (Passerine) species ^12^. “*Salmonella* virus mTmV” of strain SO4698-09, that carries the *sopE* virulence gene in some monophasic *S*. Typhimurium ST34 isolates ^18^, was absent from all other *S*. Typhimurium reference strains. However, a second prophage “*Salmonella* virus mTmII” with similarity to SJ46 was also in strain SO4698-09, and shared several clusters of gene families in common with mTmV.

With the notable exception of BcepMu the prophage predicted to be present in the common ancestor of *S*. Typhimurium exhibited considerable variation, potentially due to recombination ^31^ (Figure 4). Recombination is a major source of genetic variation in bacteria, although the level of recombination seen in other bacteria and indeed other *Salmonella* serovars vary greatly. We identified potential recombination in the genome sequence of the 131 *S*. Typhimurium of the main phylogroup by the identification of atypical SNP density. Recombination was almost exclusively present in prophage regions resulting in clade specific sequence variation (Figure 5). Recombination resulted in replacement of large blocks of gene families in ancestral prophage elements. Fels1, sal3 and Gifsy2 were conserved in the most reference strains, with the exception of Fels1 in DT104 and Gifsy2 in monophasic *S*. Typhimurium ST34 that had large alternative blocks of gene families. Gifsy1 was highly variable in all strains, but retained a core set of genes suggesting common ancestry and frequent recombination. Variation in ST64B was also present in most strains, and variable blocks of genes distinguished strains in first order clades α from β, and resulted in the acquisition of *sseK3* virulence gene by the common ancestor of the latter.

**Figure 5.**
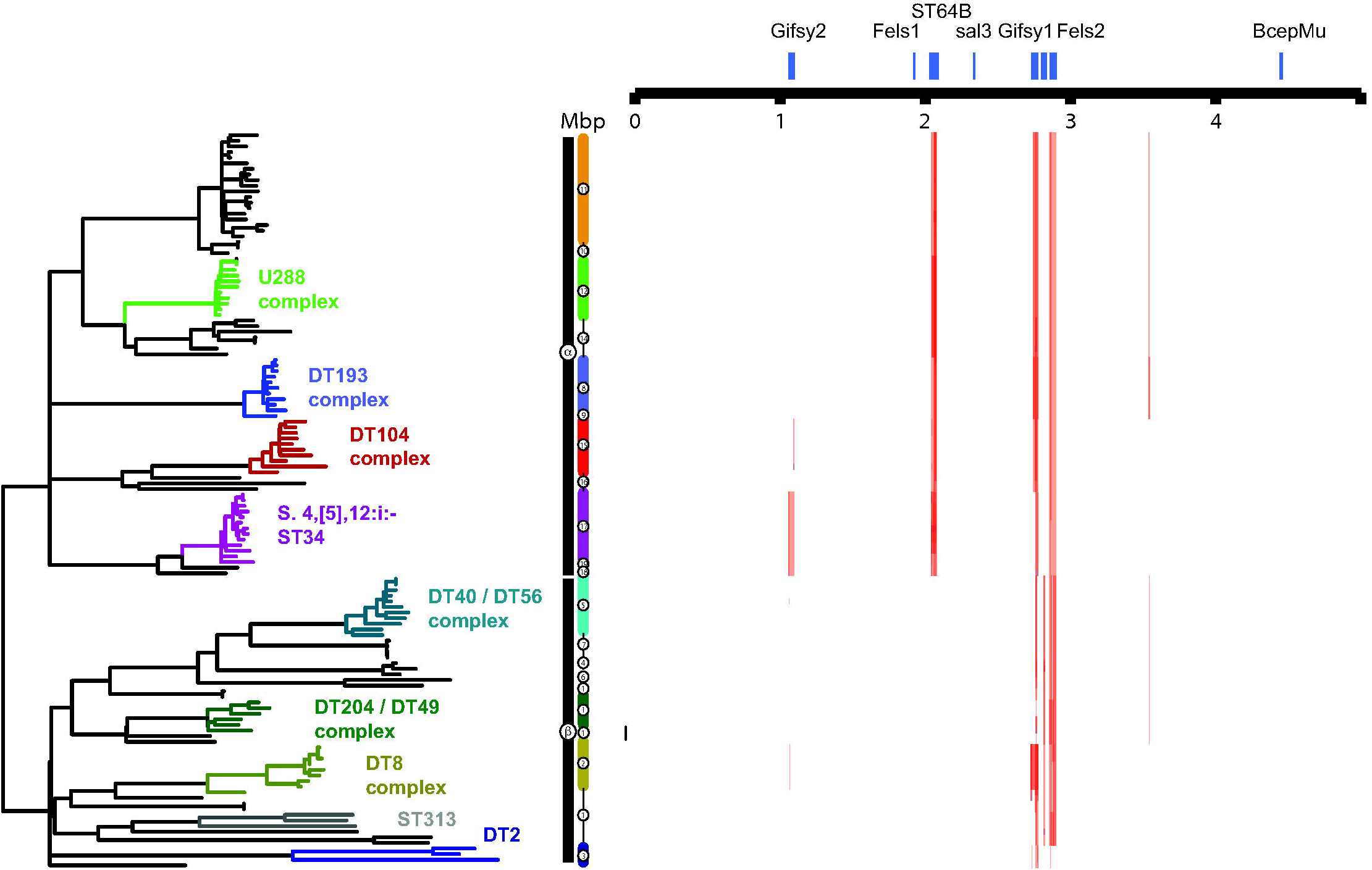
Recombination inferred by high SNP density in 131 *S*. Typhimurium strains. Regions of high SNP density (red) are indicated for each of the 131 isolates in the *S*. Typhimurium ST19 cluster with reference to the *S*. Typhimurium strain SL1344 genome. Recombination is shown with reference to the population structure and phylogeny of Typhimurium shown in figure 1. The position of predicted prophage (blue) in the *S*. Typhimurium strain SL1344 genome are indicated (top).

## Discussion

The population structure of *S*. Typhimurium consisted of two relatively distantly related clusters comprising strains of ST36 and a second (main) phylogroup the remainder of Typhimurium, but predominantly ST19, consistent with previous reports of two distinct *S*. Typhimurium phylogroups ^32, 33^. These two groups were more closely related to each other than to other serotypes of *S*. enterica subspecies I, including the nearest neighbour, serotype Heidelberg. The main Typhimurium phylogroup exhibited a star shaped phylogeny with multiple deeply rooted branches emerging from a common ancestor, with diversification at the terminal branches in some cases, associated with expansion of epidemic clonal groups. The topology of this phylogroup was similar to that of serotypes of *S*. enterica subspecies I^34, 35^, with internal branches radiating from a common ancestor, defined by the accumulation of hundreds of SNPs in *S*. Typhimurium compared with tens of thousands of SNPs defining lineages of representative strains of *S*. enterica subspecies I serotypes ^4^.

The nested phylogenetic structure, rooted with the *S*. Heidelberg outgroup, was characterised by two high order clades (α and β), in which clade α was basal to clade β. Several deeply rooted lineages of clade α contained isolates almost entirely from livestock. A single lineage originating from the common ancestor of the main *S*. Typhimurium phylogroup gave rise to the common ancestor of clade β and diversification into multiple lineages was accompanied by apparent host adaptation to diverse host species, but notably many more avian species, compared with clade α. Indeed some β subclades such as those associated with the DT56, DT2 and DT8 complexes that are well characterized host adapted clonal groups ^11, 12, 21^, contained exclusively isolates from avian species, and were present on relatively extended internal branches. This general phylogenetic topology is consistent with that described for distinct collections of *S*. Typhimurium strains from North America and Asia ^36, 37^.

Isolates in clades α and β appeared to be under distinct anthropogenic selection pressure for the acquisition and maintenance of AMR, that correlated with their distribution in livestock or wild avian species ^38^. Clade α isolates that were predominantly from livestock contained several lineages with multiple AMR genes, while most clade β isolates contained few or no AMR genes. Differential selection pressure for acquisition and maintenance of AMR genes is consistent with the idea that some *S*. Typhimurium genotypic variants are adapted to circulation in specific host populations that exert different selection pressure for the acquisition and maintenance of AMR genes. Antimicrobials have been used widely to control infection or as growth promoters in livestock, but wild animals are unlikely to encounter therapeutic levels of these drugs ^39^. However, *S*. Typhimurium strains of DT56 and DT40 that we report are present in clade β, known to be associated with passerine birds, and lacking AMR genes have also been reported in cattle where they may be subject to selection for antimicrobial resistance ^40^, and we might therefore expect to find AMR genes in these strains also. One possibility is that DT56 and DT40 strains from clade β may transiently colonise the cattle host, but are unable to circulate in this population and do not transmit back to the avian population with high frequency. Host adaptation to avian species therefore appears to create an effective barrier to circulation in livestock. Two clade β lineages did contain strains with multiple AMR genes, but in each case, they were atypical for this clade in that they were associated with an epidemic in cattle (DT204/49 complex) or invasive NTS in people in sub-Saharan Africa (ST313) ^7, 22^, and therefore were likely to be under selection for AMR.

The molecular basis of the barrier to circulation of several clade of the β isolates in livestock is not known, but likely the result of genotypic changes affecting functional diversification of the proteome. The proteome delta bitscore (DBS) of clade β isolates exhibited elevated divergence from profile HMMs of protein families in gamma proteobacteria, compared to clade α isolates, potentially resulting in loss or altered protein function ^21^. Similarly, divergence was reported in the *S*. Gallinarum proteome, a serotype highly host adapted to poultry where it is associated with fowl typhoid ^25^. β subclades also exhibited an elevated invasiveness index, the fraction of decision trees using random forests that vote for the invasive (extraintestinal) disease outcome, a predictive score of host adaptation to an extraintestinal lifestyle. In our analysis DBS of 300 protein families most predictive of disseminated disease in *S*. Typhi, *S*. Patatyphi A, *S*. Gallinarum, *S*. Dublin and *S*. Choleraesuis, also predicted an extraintestinal lifestyle in *S*. Typhimurium β subclades, consistent with the epidemiological data ^4^. The pattern of functional divergence in some *S*. Typhimurium β subclades may therefore at least in part be by a process of convergent evolution with that observed as a result of the evolution of several extraintestinal serotypes *S. enterica*, including *S*. Typhi, *S*. Patatyphi A, *S*. Gallinarum, *S*. Dublin and *S*. Choleraesuis ^26^.

Host adaptation of bacteria is commonly associated with the accumulation of HDCS, potential pseudogenes that contribute to genome degradation ^41, 42^. A total of 24 genes previously implicated in virulence, adhesion or multicellular behaviour were HDCS in one or more clade β isolates. Notably, 15 of these genes (63%) were also HDCS in highly host adapted *S*. Typhi or *S*. Paratyphi A, serotypes that are restricted to humans and cause a disseminated disease^43^. HDCS were especially common in strains of DT56/DT40 complex (clade β5), that are reported to be highly host adapted to passerine birds, in which eleven HDCS were observed. In *S*. Paratyphi A, many mutations and gene flux that occurred was reported to be neutral, indicated by their sporadic distribution within clades, and frequent loss from the population by purifying selection ^44^. In *S*. Typhimurium, we also observed some examples of potential neutral mutations, but in many cases HDCS in virulence genes were present in multiple related strains from the same subclade, indicating that they were likely under selection, and stably maintained in the population (Figure 1). In contrast to clade β, genome degradation affecting virulence-associated genes was less frequent in isolates clade α. Just three virulence gene HDCS had a clade phylogenetic signature, *avrA*, *tsr* and *sadA*, in α10, α11 and α12. Subclade α12 (U288 complex) was the only clade to contain all of these HDCS, consistent with the U288 complex exhibiting apparent host-adaption to pigs ^17, 29^. Therefore, the presence of HDCS in virulence associated genes was almost exclusively associated with subclades containing strains with strong epidemiological evidence of host adaptation ^4^. Despite the *S*. Typhimurium DT104 complex strain NCTC13348 (clade α15) exhibiting a high level of genome degradation that was uncharacteristic for α subclades and the broad host range epidemiology of the clonal group ^45^, no virulence or multicellular behaviour genes were HDCS ^45^. Furthermore, the mean DBS for the proteome of strains from clade α15 was similar to that of other α subclades, suggesting that functional divergence as a whole was not atypical from that of other clade α isolates.

While genes encoding components of the type III secretion systems (T3SS) 1 and 2 apparatus were never HDCS, several genes encoding effector proteins secreted by them were (*sseI*, *sseK2, sseK3*, *avrA*, *sseL*, *sseJ* and *gtgE*). The *sseI* gene is inactivated in ST313 due to insertion of a transposable element, and results in hyper dissemination of these strains to systemic sites of the host via CD11b^+^ migratory dendritic cells ^46^. The *sseK2, sseK3*, *avrA* and *sseL* genes each encode effectors that inhibit the NFκB signalling pathway thereby modulating the proinflammatory response during infection ^47–49^. Furthermore, these effectors are commonly absent or degraded in serotypes of *Salmonella* serotypes associated with disseminated disease ^50, 51^, suggesting that altered interaction with the macrophage is essential for disseminated disease in diverse *Salmonella* variants and hosts. In addition, several genes encoding components of fimbrial or afimbrial adhesin systems (*sadA*, *ratB*, *lpfD*, *stfD*, *stbD*, *safC*, *fimH*, *siiC* and *siiE*), or anaerobic respiration (*ttrB*), and chemotaxis (*tsr*) were HDCS in one or more isolates. Many of these genes have been implicated in intestinal colonisation^52–58^, suggesting that their inactivation in host adapted variants may be associated with a loss of selection for functions no longer required in a reduced host range or where intestinal colonisation is no longer critical to transmission. The *sadA* and *katE*, genes are involved in multicellular behaviour, biofilm formation and catalase activity that protects against oxidative stress during high density growth functions, respectively, and are commonly affected by genome degradation in host adapted pathovars of *Salmonella* ^59, 60^.

The only virulence-related HDCS that segregated clades α and β resulted from a 10 bp insertion in the *lpfD* gene of all clade β isolates. Within the host *S*. Typhimurium preferentially colonises Peyer’s patches (PPs) ^61^, due to long-polar fimbriae *Lpf* binding to M-cells ^62^. Similarly, *lpf* genes in *E. coli* pathotypes are required for interaction with Peyer’s patches and intestinal colonisation ^63^. Despite the disruption of *lpfD* in the clade β isolate SL1344, deletion of *lpf* reduced colonisation on the surface of chicken intestinal tissue, ^64^, suggesting that long polar fimbriae retain function. Differences in intestinal architecture of avian species that have the lymphoid organ the bursa of Fabricius containing numerous M cells compared with mammalian species that have Peyer’s patches with relatively scarce M cells ^65, 66^ may explain the pattern of *lpfD* HDCS in S. Typhimurium. The function of long polar fimbriae expressing full length LpfD in clade α isolates has not been investigated, but its distribution in isolates from livestock and human infections mark it as of potential importance to human health.

Bacterial genome diversity is largely driven by the flux of genes resulting from acquisition by horizontal gene transfer and deletion, rather than allelic variation ^67^. The accessory genome of *S*. Typhimurium revealed few genes that segregated clade α and β, but distinct forms of ST64B prophage resulting from recombination that replaced a large block of genes were present in clade α and β, and resulted in the presence of *sseK3* specifically in clade β. The accessory genome contributed significantly to genetic variation that distinguished third order subclades in both clade α and β, especially phage and plasmid genes. Non-phage chromosomal genes exhibited relatively little clade specific accessory genome suggesting that the majority was the result of deletions or gene acquisition on small mobile genetic elements that were neutral and subsequently lost, as observed previously in *S*. Paratyphi ^44^. However, three large genetic elements were acquired on the chromosome in α15 (DT104) or α17 (monophasic *S*. Typhimurium ST34), the two most recent dominant MDR pandemic clonal groups that together account for over half of all S. Typhimurium infections in the human population Europe in the past 30 years. The acquired genes corresponded to SGI1 ^28^ in the DT104 complex, and SGI4 and a composite transposon in monophasic *S*. Typhimurium ST34 ^18, 29^, highlighting the likely importance of horizontal gene transfer in the emergence of epidemic clones.

Variable prophage repertoires are a major source of genetic diversity in *Salmonella* ^68, 69^, and may contribute to the emergence and spread by impacting the fitness during intra-niche competition due to lytic killing or lysogenic conversion of competing strains ^70^. This view was supported by the considerable phage-associated gene flux observed in *S*. Typhimurium. Importantly, the phage component of the accessory genome in *S*. Typhimurium had a strong correlation with the third level clade, suggesting that although transfer was frequent, the acquisition or loss of prophage elements was not transient, consistent with selection within each clonal group ^44^. Reconstruction of the evolutionary history of prophage elements in the main *S*. Typhimurium phylogroup indicated that the common ancestor likely contained six prophage, Gifsy1, Gifsy2, Fels1, ST64B, sal3 and BcepMu, that were well conserved during subsequent diversification. Just two of these ancestral prophage, Fels1 and Gifsy1, were lost from the genome, on three occasions in different lineages. The majority of the prophage flux was from the acquisition of between one to three prophage in each lineage, with the exception of a lineage containing clade β3 (DT2), that only contained the ancestral prophage repertoire.

Together, our analyses are consistent with the view that the common ancestor of the main *S*. Typhimurium phylogroup was a broad host range pathogen with little genome degradation capable of circulating within multiple species of livestock. The age of the common ancestor of *S*. Typhimurium is not known and previous attempts to calculate this using Bayesian approaches have been frustrated by a weak molecular clock signal ^71^. However, the common ancestor of *S*. Paratyphi A was estimated to have existed approximately 500 years ago and provides a frame of reference ^44^. The main *S*. Typhimurium phylogroup exhibited greater genetic diversity than reported for *S*. Paratyphi A, an estimated maximum root to tip SNP accumulation of approximately 750 and 250, respectively. Therefore, the common ancestor of the main *S*. Typhimurium phylogroup is likely to have existed substantially before that of *S*. Paratyphi A. However, the S. Typhimurium MRCA is unlikely to have predated the domestication of livestock, that began around ten thousand years ago ^72^, raising the possibility that the emergence of this phylogroup was linked to the anthropogenic selection provided by the domestication of species for livestock. Subsequent to the emergence of the common ancestor of this phylogroup, a single lineage appears to have spawned multiple lineages, some of which have become highly host adapted to various wild avian species, by a process of convergent evolution with that observed in host adapted serotypes of *Salmonella* such as *S*. Typhi.

## Materials and Methods

### Bacterial strains and culture

*S*. Typhimurium isolates and Illumina short read sequence used in this study have been described previously ^73^, selected based on phage type determined during routine surveillance by Public Health England (PHE) and the Animal and Plant Health Agency (APHA) in order to represent the diversity *S*. Typhimurium phage types as a proxy for genetic diversity. A strain collection of 134 *S*. Typhimurium or monophasic variant isolates was composed of 2 to 6 randomly selected strains from the top ten most frequent phage types from PHE and the top 20 most frequent phage types from APHA surveillance, from 1990-2010 (2-5 strains of each) were used in this analysis. In addition, commonly used lab strain SL1344 ^74^, two reference strains of ST313 (D23580 and A130)^75^ and three DT2 strains isolated from pigeon ^76^ were included. For routine culture, bacteria were stored in 25% glycerol at -80°c and recovered by culture on Luria Bertani agar plates, and single colonies were selected to inoculate LB broth that was incubated at 37 °c for 18 hours with shaking.

### Short-read *de novo* assembly

Illumina generated fastq files were assembled using an in-house pipeline adapted from that previously described ^77^. For each paired end reads, Velvet ^78^ (1.2.08) was used to generate multiple assemblies varying the k-mer size between 31 and 61 using Velvet Optimiser ^79^ and selecting the assembly with the longest N50. Assemblies were then improved using Improve_Assembly software ^80^ that uses SSPACE (version 3.0) ^81^ and GapFiller (version 1.0) ^82^ to scaffold and gap-fill. Ragout was used to order contigs ^83^ based on comparison to the long-read sequences. The finished genomes were then annotated using Prokka (version 1.11) ^84^.

### Long-Read sequencing using Pacbio and sequence assembly

DNA for long-read sequencing on the Pacbio platform was extracted from 10 ml of cultured bacteria as previously described ^73^. Data were assembled using version 2.3 of the Pacbio SMRT analysis pipeline (https://smrt-analysis.readthedocs.io/en/latest/SMRT-Pipe-Reference-Guide-v2.2.0/). The structure of the initial assembly was checked against a parallel assembly using Miniasm ^85^ which showed general agreement. The Pacbio best practice for circularizing contigs was followed using Minimus ^86^ and the chromosomal contiguous sequence in each assembly was re-orientated to begin at the *thrA* gene. Illumina short read sequence data were used to correct for SNPs and indels using iCORN2 (http://icorn.sourceforge.net/). The finished sequences were then annotated using Prokka ^84^.

### Phylogenetic reconstruction and population structure analysis

The paired-end sequence files for each strain were mapped to the SL1344 reference genome (FQ312003) ^74^ using the Rapid haploid variant calling and core SNP phylogeny pipeline SNIPPY (version 3.0) (https://github.com/tseemann/snippy). The size of the core genome was determined using snp-sites (version 2.3.3) ^87^, outputting monomorphic as well as variant sites and only sites containing A,C,T or G. Variant sites were identified and a core genome variation multifasta alignment generated. The core genome of 134 *S*. Typhimurium (3686476 nucleotides) (Supplementary Table 2) contained 17823 variant sites. The core genome (3739972 nucleotides) of 131 *S*. Typhimurium non-ST36 contained 8382 variant sites. The sequence alignment of variant sites was used to generate a maximum likelihood phylogenetic tree with RAxML using the GTRCAT model implemented with an extended majority-rule consensus tree criterion ^88^. The genome sequence of *S.* Heidelberg (NC_011083.1) was used as an outgroup in the analysis to identify the root and common ancestor of all *S*. Typhimurium strains. HierBaps (hierarchical Bayesian analysis of Population Structure) ^16^ was used to estimate population structure using three nested levels of molecular variation and 10 independent runs of the optimization algorithm as reported previously ^89^. The input for this analysis was the same SNP variant matrix for the 131 strains with reference to SL1344 that was used to generate the GTRCAT phylogeny above.

### *In-silico* genotyping

The presence of antibiotic resistance, virulence and plasmid replicon genes in short-read data was determined by the mapping and local assembly of short reads to data-bases of candidate genes using Ariba ^90^. The presence of candidate genes from the resfinder ^91^, VFDB ^92^ and PlasmidFinder ^93^ databases was determined. Reads were mapped to candidate genes using nucmer with a 90% minimum alignment identity. This tool was also used to determine the presence of specific genes or gene allelic variants. The results of the ARIBA determination of the presence or absence of the *lpfD* gene were confirmed using SRST2 ^94^ setting each alternative form of the gene as a potential allele. SRST2 was also used to verify the ARIBA findings of the VFDB data set, as the presence of orthologous genes in the genome was found to confound the interpretation of results.

### Hypothetically disrupted coding sequences (HDCS)

HDCS were identified in high-quality finished Pacbio sequences and previously published reference sequences by identifying putative altered open reading frames using the RATT annotation transfer tool ^95^. The *S.* Typhimurium strains SL1344 annotation (accession no. FQ312003) was transferred to each assembled sequence and coding sequences identified as having altered length were manually curated by comparison of aligned sequences visualised using Artemis comparison tool (ACT)^96^. Genes that contained either a premature stop codon or a frameshift mutation were classified as HDCS. The identified HDCS were used to construct a database that could be used as a reference for SRST2 (above) to detect presence or absence in short-read sequence data. Alleles were called based on matching to 99% sequence identity and allowing one miss-match per 1000 nucleotides.

### Delta bitscore

Illumina short-read sequences were mapped to the SL1344 reference genome and annotated using PROKKA and then analysed in a pairwise fashion against SL1344 using delta-bit-score (DBS), a profile hidden Markov model based approach ^25^ with Pfam hidden Markov Models (HMMs) ^97^. The mean DBS per genome and percentage of genes with mutations in Pfam domains (non-zero DBS) are reported.

### Invasiveness Index

The invasiveness index ^26^ for each strain was calculated to scan for patterns of mutation accumulation common to *Salmonella* lineages adapted to an invasive lifestyle. To calculate the invasiveness index, Illumina reads were mapped to a core-genome reference using the snippy pipeline above, and annotated using PROKKA. Protein sequences were then screened using phmmer from the HMMER3.0 package ^98^ to identify the closest homologs to the 196 predictive genes used by the invasiveness index model. These genes were then scored against profile hidden Markov models (HMMs) for these protein families from the eggNOG database ^26^ using hmmsearch ^98^, to test for uncharacteristic patterns of sequence variation. Bitscores produced in the comparison of each protein sequence to its respective protein family HMM were then used as input to the model.

### Phage location and cargo in long-read strains

The location of prophage elements in assembled long-read sequences and published reference genome was determined using Phaster ^99^, which identified regions as being intact, questionable or incomplete. This yielded a total of 83 potential complete and partial sequences across the 11 representative strains. Prophage sequences were annotated using PROKKA that identified terminase, tail fibre, recombinase/integrase proteins capsid proteins, phage related proteins, and hypothetical proteins.

### Determination of recombination

Recombination was inferred by identifying regions of high SNP density from whole genome alignments of short-read data to SL1344, using Gubbins ^100^. The results were visualised using Phandango ^101^ and related to the predicted prophage locations in the SL1344 genome. Similar results were obtained using maximum likelihood inference using clonal frame ^102^.

### Determination of the *S*. Typhimurium pangenome

The annotated assemblies of 131 predominantly *S*. Typhimurium ST19 isolates were used as the input to the pangenome pipeline ROARY ^103^. The presence or absence of genes was determined without splitting orthologues. In order to characterise the contribution of prophage and plasmids to the pangenome, genes were assigned to one of four categories, non-prophage genes located on the chromosome, prophage genes, plasmid genes and undefined genes, based on the similarity to annotated genes of complete and closed whole genome sequence of eleven reference strains. Orthologous genes were identified based on > 90% nucleotide sequence identity using nucmer ^104^. A core-genome reference sequence (genes present in at least 99% of reference strains), was also constructed and used to determine the invasiveness index.

### Estimation of gene flux rates

Genes were assigned a score based on their presence in strains within a specific clade. The clade gene score was compared to the score determined for strains outside of the clade to determine whether the gene was more prevalent within the clade than without. Genes were classed as associated with the clade if their score was greater than the mean plus two standard deviations of the non-cladal score (corresponding to the top 95% in a normal distribution).

The number of clade associate genes was compared with the number of SNPs associated with a clade (this gives a measure of evolutionary time) to determine the level of gene flux for the clade. The level of gene flux in the two first-level clades was then compared.

### Prophage classification

PHASTER^99^ curated prophage sequences were classified into species and genus-level groupings based on the current criteria used by the Bacterial and Archaeal Viruses Subcommittee of the International Committee on Taxonomy of Viruses (ICTV)^105^. At the species level, genomes were clustered at 95% nucleotide identity over the whole genome length, meaning that two genomes belong to two different species if they differ in more than 5% of their genome. Clustering was performed with CD-HIT-EST at 95% nucleotide identity over 95% of the alignment length (99% of alignment length of shorter sequence)^106^ and with Gegenees, a pairwise nucleotide comparison tool, using the accurate settings of 200 bp fragment size and 100 bp step size^107^. The Gegenees output was used in combination with vConTACT2^108^ to classify the prophage sequences into new or existing genera. Briefly, coding sequences were predicted with PROKKA^84^ and transformed into a table linking genomes and their encoding proteins. This table was used as input into vConTACT along with the Viral RefSeq database v85^109^. vConTACT then used Diamond^110^, Markov clustering MCL^111^ and ClusterONE^112^ to predict viral clusters based on shared protein content. The output was visualised using Cytoscape^113^. Genera were defined as vConTACT viral clusters which shared a significant (>50%) nucleotide identity. The clusters were then compared to the current and pending ICTV taxonomic classification (ictvonline.org) using blast and vConTACT viral cluster output containing reference genomes and all prophage sequences were assigned to new or existing taxa.

## Declarations and Acknowledgements

No procedures or data collection required ethics approval, consent to participate, or consent for publication. All data is available in accessible databases as indicated in the text and supplementary information tables. The authors declare that they have no competing interests. The project was conceived by MB and RAK, data analysis was performed by MB, GT, MK, NW and EMA, interpretation of the data was by MB, RAK, LP, TD and NH, the manuscript was drafted by MB and RAK, and edited and approved by GT, MK, NW, LP, TJD, EMA, NH and RAK. The author(s) gratefully acknowledge the support of the Biotechnology and Biological Sciences Research Council (BBSRC); RAK was funded by the BBSRC Institute Strategic Programme Microbes in the Food Chain BB/R012504/1 and its constituent project(s) BBS/E/F/000PR10348 and BBS/E/F/000PR10352, and by projects BB/J004529/1, BB/M025489/1 and BB/N007964/1. NH was supported by a BBSRC funding for the Earlham Institute BB/CCG1720/1. EMA was supported by the BBSRC Institute Strategic Programme Gut Microbes and Health BB/R012490/1 and its constituent project BBS/E/F/000PR10356. The genome sequencing for this work was carried out by the Genomics Pipelines group at the Earlham Institute which is funded as a BBSRC National Capability (BB/CCG1720/1). EMA was funded by the BBSRC Institute Strategic Programme Gut Microbes & Health BB/R012490/1 and its constituent projects BBS/E/F/000PR10353 and BBS/E/F/000PR10356. This research was supported in part by the NBI Computing infrastructure for Science (CiS) group through use of HPC resources. The authors also acknowledge advice and informatics support from Andrew Page, Andrea Telatin and Nabil-Fareed Alikhan from the Quadram Institute Bioscience informatics support group.

---

**Supplementary Figure 1. Phylogenetic relationship of *S*. Typhimurium and diverse *S*. enterica serotypes.** Mid-point rooted maximum likelihood phylogenetic tree based on the variation (SNPs) in the core genome of 18 strains of *Salmonella* Typhimurium and 14 representative strain of diverse *S*. *enterica* subspecies *enterica* serotypes, with reference to *S*. Typhimurium strain SL1344 genome sequence. *S*. Typhimurium strains (red lineages and text) are present in two clusters, composed of 15 strains with isolates that are ST19, ST34, ST313, ST98 and ST568 and three more divergent isolates of ST36. The phylogeny is rooted with respect to *S.* Heidelberg and was calculated using SL1344 as a reference to create a core-genome variant-site alignment and the GTRCAT model in RAxML.

**Supplementary figure 2. Accessory genome with strong clade association**. Gene families with a strong clade association in clade a or b (A), or in one of the third level clades (B). Maximum likelihood phylogenetic tree and based on sequence variation (SNPs) in the core genome with reference to *S*. Typhimurium strain SL1344 (left). Third-level clades are indicated and colour coordinated with that in Figure 1. Genes in each clade were assigned a score based on the number of strains containing the gene within the clade. This score was also calculated for the strains outside the clade. Clade associated genes were defined as genes that had scores greater than the mean plus two SD of the score for all other clades. Genes are colour coded based assignment to non-prophage chromosomal (red), prophage (green), plasmid (blue), or undefined (grey).

**Supplementary Figure 3.** Gene flux rate metrics determined for non-singleton gene families first-level clades.

## Supplementary Table Legends

**Supplementary Table 1. S. Typhimurium strain collection used in this study related to determine population structures**. Table can be viewed at https://www.dropbox.com/sh/dh84yyc4tguirw3/AABnbbrSPtqEcjGY6IR6GggLa?dl=0

**Supplementary Table 2.** Presence of plasmid operons, AMR genes (Resfinder) and virulence genes (VFDB) determined by *in-silico* genotyping using ARIBA software. The identifier column corresponds to the study-identifier column in Supplementary Table 1. Presence of genes are indicated by ‘1’. Table can be viewed at https://www.dropbox.com/sh/dh84yyc4tguirw3/AABnbbrSPtqEcjGY6IR6GggLa?dl=0

**Supplementary Table 3. Genome degradation in long-read reference strains.** Potential hypothetically disrupted coding sequences (HDCS) were identified in reference genomes though anomalies in annotation transfer from the SL1344 reference sequence using RATT software and manual curation to exclude false positive HDCS. Table can be viewed at https://www.dropbox.com/sh/dh84yyc4tguirw3/AABnbbrSPtqEcjGY6IR6GggLa?dl=0

**Supplementary Table 4. Summary of HDCS in S. Typhimurium main phylogroup.** The presence of HDCS alleles was determined *in-silico* using SRST2 reported in Figure 1. The study identifier refers to isolates in Supplementary Table 1. Alleles are specified as wild-type (WT) or HDCS. In some cases, multiple HDCS forms were determined to be present and these are denoted as HDCS1 or HDCS2 etc. Table can be viewed at https://www.dropbox.com/sh/dh84yyc4tguirw3/AABnbbrSPtqEcjGY6IR6GggLa?dl=0

**Supplementary Table 5. Summary of DBS and Invasiveness Index analysis.** Isolate identifier corresponds to the identifier column in Supplementary Table 1. The mean delta bitscore (DBS) for the proteome of each strain, the number of proteins with a DBS greater than ten. Invasiveness index for each isolate is indicated. Table can be viewed at https://www.dropbox.com/sh/dh84yyc4tguirw3/AABnbbrSPtqEcjGY6IR6GggLa?dl=0

**Supplementary Table 6. Characteristics of prophage elements present in complete and closed whole genome sequence of S. Typhimurium reference strains**. Table can be viewed at https://www.dropbox.com/sh/dh84yyc4tguirw3/AABnbbrSPtqEcjGY6IR6GggLa?dl=0

