## Supplementary figures and images for "Evolution of *Salmonella enterica* serotype Typhimurium driven by anthropogenic selection and niche adaptation"

### Supplementary Figure 1

Supplementary Figure 1

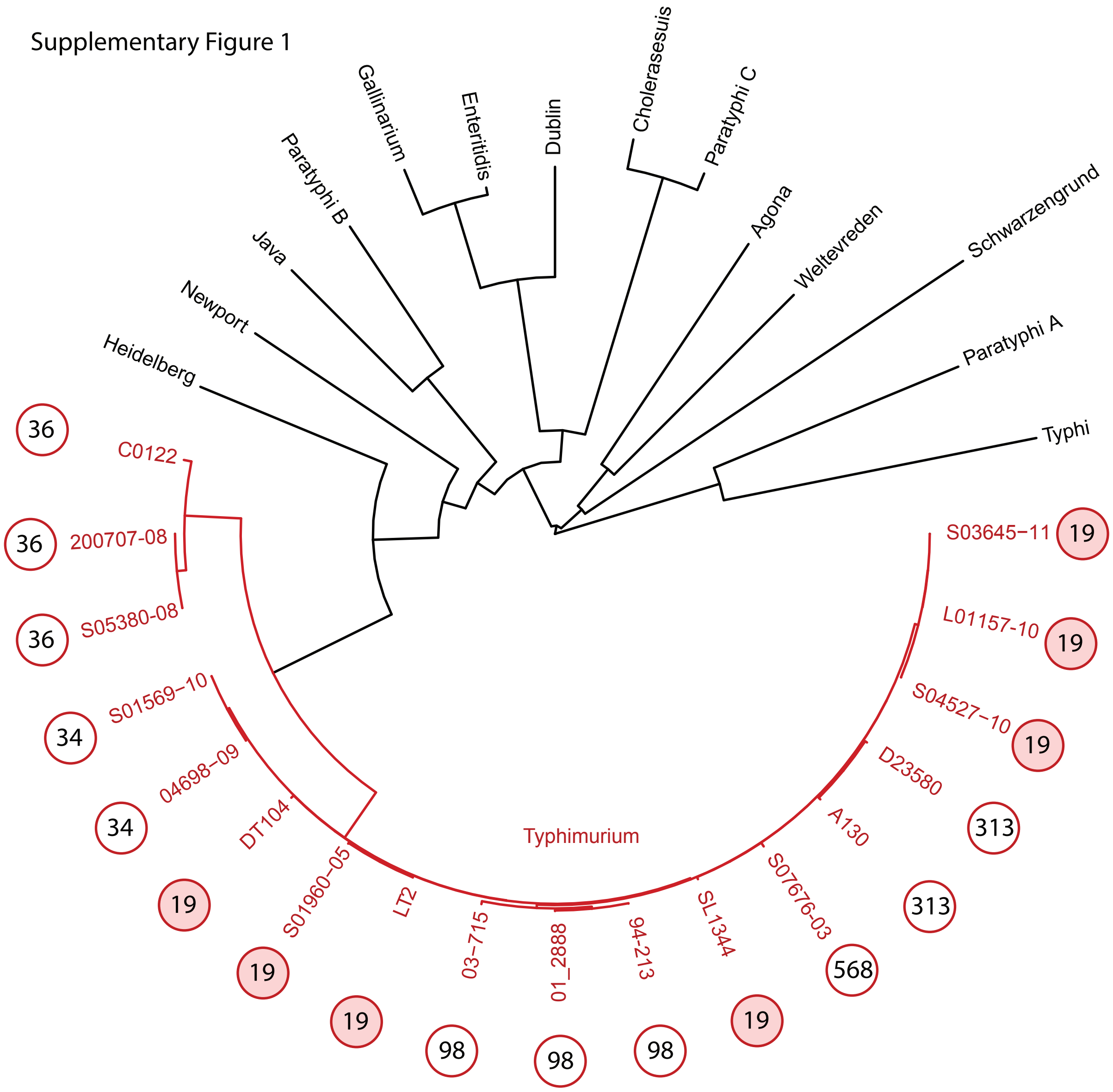

### Supplementary Figure 2

Supplementary Figure 2

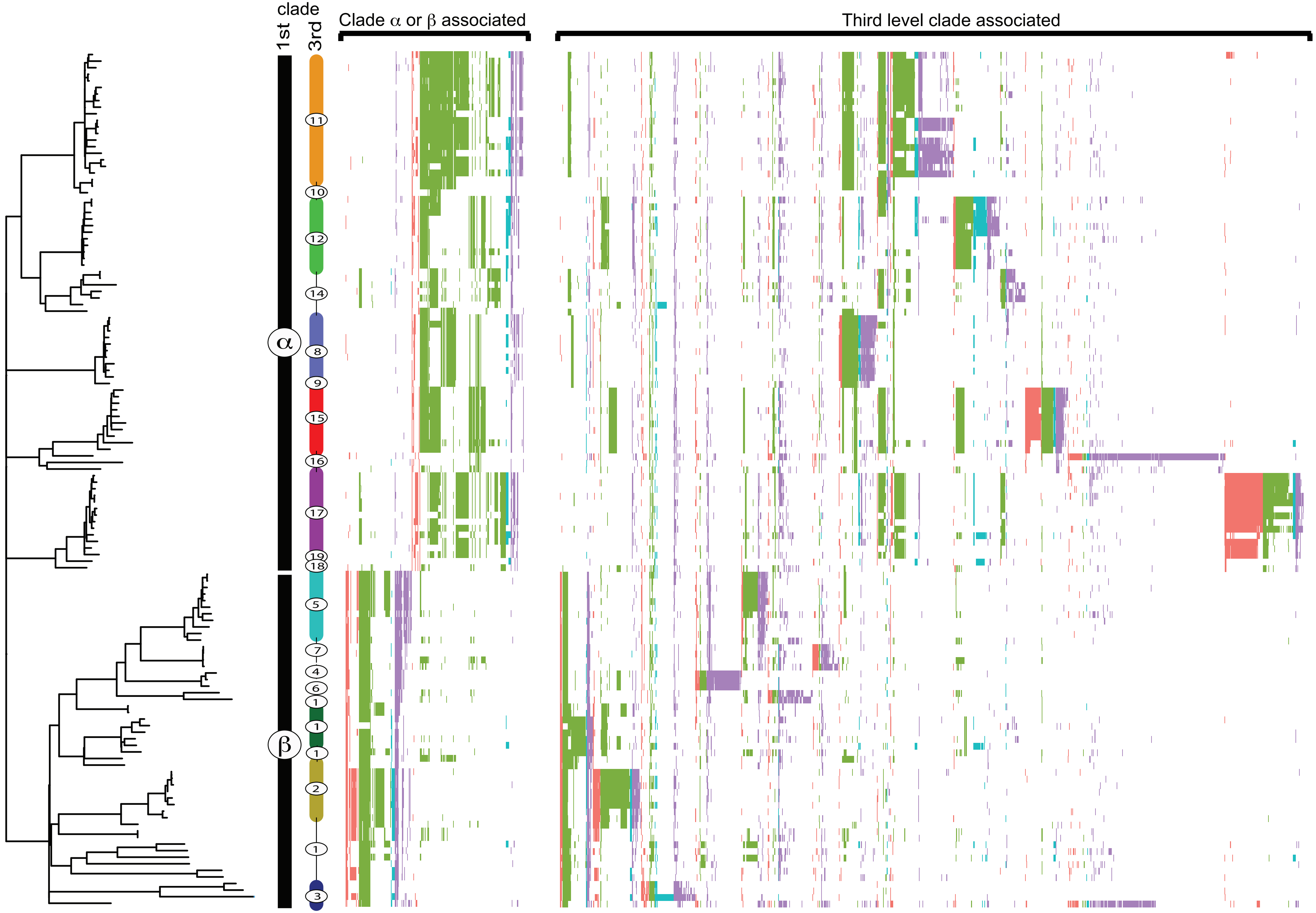

### Supplementary Figure 3

Supplementary Figure 3

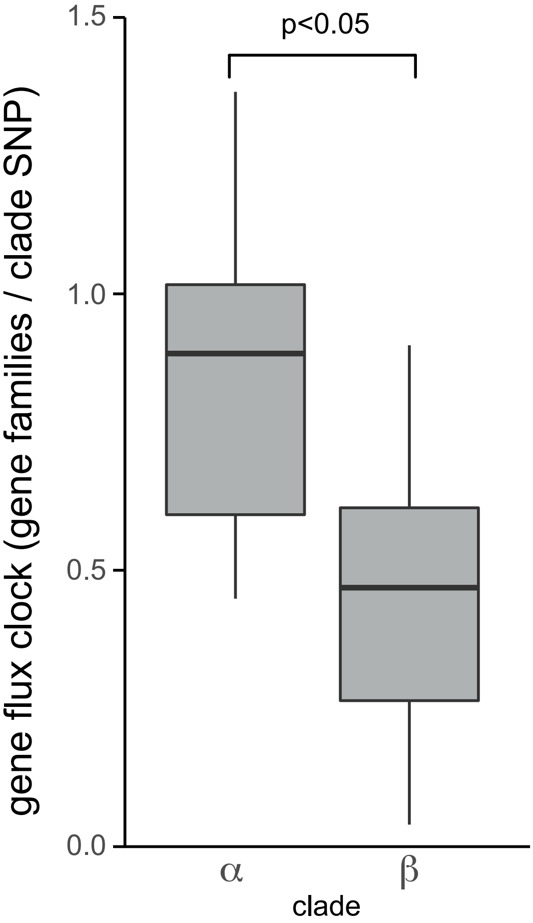
